# Utility of FRET in studies of membrane protein oligomerization: the concept of the effective dissociation constant

**DOI:** 10.1101/2023.05.16.540587

**Authors:** Daniel McKenzie, Daniel Wirth, Taras V. Pogorelov, Kalina Hristova

## Abstract

The activity of many membrane receptors is controlled through their lateral association into dimers or higher order oligomers. While Förster resonance energy transfer (FRET) measurements have been used extensively to characterize the stability of receptor dimers, the utility of FRET in studies of larger oligomers is unclear. Here we show that we can extract an effective equilibrium dissociation constant from FRET measurements for EphA2, a receptor tyrosine kinase (RTK) known to form active oligomers of heterogeneous distributions in response to its ligand ephrinA1-Fc. The newly introduced effective equilibrium dissociation constant has a well-defined physical meaning and biological significance. It denotes the receptor concentration for which half of the receptors are monomeric and inactive, and the other half are associated into oligomers and are active, irrespective of the exact oligomer size. This work illustrates how FRET, along with fluorescence fluctuation techniques which directly measure the oligomer size, can be a very powerful tool in studies of membrane receptor association and signaling in the plasma membrane.

## INTRODUCTION

Membrane proteins are abundant in eukaryotes, and account for 20% to 30% of the open reading frames(1). They play key roles in cell signaling, cell adhesion, recognition, motility, energy production, and transport of nutrients. Just as in the case of soluble proteins, the function of membrane proteins is often regulated through their homointeractions or through heterointeractions with partner proteins(2-7). However, while soluble proteins are typically studied in purified form using well established quantitative methods, membrane proteins are fickle and easily lose their activity once extracted out of the native membranes they reside in(8). Thus, often the only viable option is to study them in the context of the complex membrane milieu (9, 10). Because of such restrictions, knowledge of the folding, structure, and function of membrane proteins has been slow to emerge.

Studies of the second largest class of membrane receptors, the RTKs, exemplify challenges and limitations in membrane protein research. RTKs are single-pass transmembrane proteins that control cell growth, differentiation, motility, and metabolism (11, 12). They play profound roles in human development and are strongly implicated in disease. Their N-terminal extracellular (EC) regions, composed of characteristic arrays of structural domains, bind the activating ligands(13). They have single transmembrane helices and intracellular kinase domains. Despite their simple architecture and their significance for human health, there are currently no high-resolution structures of any of the 58 full-length RTKs, and there is no detailed mechanistic understanding of their activation(14). What is well known is that the function of RTKs is regulated through their self-association in the membrane. RTK monomers are inactive while RTK dimers/oligomers are active, as kinases in close proximity cross-phosphorylate each other by acting as both enzymes and substrates(5, 15-18). This cross-phosphorylation stimulates catalytic activity, resulting in the phosphorylation of cytoplasmic substrates and downstream signaling(5, 19-21).

For many years, the activation of RTKs was believed to follow the simple model of ligand induced dimerization(13, 22). Later, it was shown that many RTKs dimerize even in the absence of ligand, and that ligand binding stabilizes these dimers(6, 23). Dissociation constants in the absence and in the presence of high saturating ligand concentrations have been measured using FRET(24-28). In these experiments, RTK expressions are varied over a wide range, and RTK concentrations in the plasma membrane are measured in hundreds of cells, along with FRET efficiencies, yielding dimerization curves and dissociation constants.

Recent work has revealed that some RTKs signal as oligomers that are larger than dimers(29-32). In such cases, interpretation of FRET data is more challenging(33). Here we introduce the concept of the “effective dissociation constant”, which allows us to understand and predict the self-association, and therefore the activities, of RTKs even when their association state is unknown. We assess the utility of the concept using the receptor EphA2, which plays an important role in cell guidance during development and has been implicated in many cancers. EphA2 oligomerization has been characterized using fluorescence fluctuation methods such as PIE-FCCS, N&B and FIF(29, 34, 35). These methods can directly report on the distribution of oligomer sizes, but have had limited use in determining oligomer stability. Prior fluorescence fluctuation studies have revealed that the size distributions of the ligand-bound EphA2 oligomers is heterogeneous, with average oligomer size of ∼4(36). In this work we measure FRET for EphA2 in the presence of its ligand ephrinA1-Fc, and we calculate the effective dissociation constant.

## RESULTS

### Theory: The Effective Dissociation Constant

#### The dissociation constant describing protein-ligand interactions

Equilibrium constants describe the relative abundances of reactants and products for a reaction at equilibrium and report on which are favored. One such reaction is the binding of a ligand, *L*, to a protein, *P*, to form a complex, *LP*.

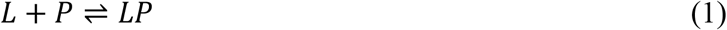

The association equilibrium constants *K*_*A*_ is defined as:

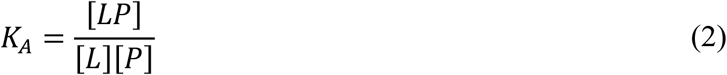

where [*L*] denotes the free ligand concentration, [*P*] is the free protein concentration, and [*LP*] is the concentration of the complex. The dissociation constant, *K*_*D*_, is the inverse of *K*_*A*_:

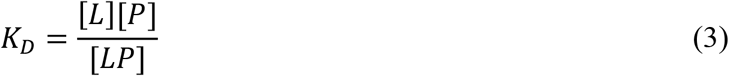

The units for each constant are important to note, with *K*_*A*_ being inverse concentration and *K*_*D*_ being concentration. Of the two equilibrium constants, the usage of *K*_*D*_ is preferred because *K*_*D*_ has units of concentration, and its value can be compared to the reactants concentrations. At low reactant concentrations, the free reactants are favored. A high reactant concentrations, higher than *K*_*D*_, the products are favored.

The dissociation constant has another feature, namely if [*LP*] = [*P*], then *K*_*D*_ = [*L*]. Thus, *K*_*D*_ is the free ligand concentration at which the fraction of bound protein is 50% (0.5). Often, *K*_*D*_ is approximated as the total ligand concentration for which the bound protein fraction is 50%. This approximation is made when the free ligand concentration is unknown, but is not always valid, depending on the experimental design.

#### Dissociation constants describing receptor self-association

The simplest case of homointeractions is dimerization, described by the following reaction scheme:

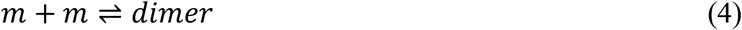

where *m* denotes the monomer. The dissociation constant is:

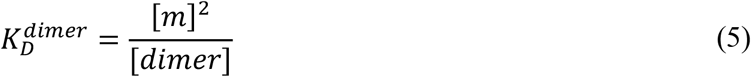

Brackets indicate the concentrations of monomers and dimers. *K*_*D*_ in reaction (4) has similar biological significance as in reaction (1) and has the same units, concentration. However, since the interactions occur in the two-dimensional membrane, the units are receptors per unit area. The fraction of dimeric receptors is given by:

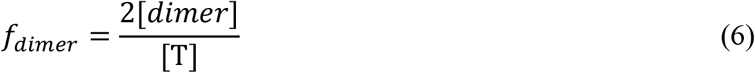

Where [T] is the total receptor concentration, [*m*] + 2[*dimer*]. Substitution of equation (5) into equation (6) yields:

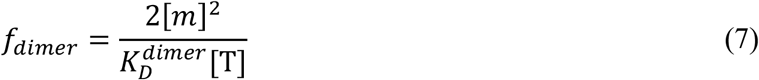

Now we write equation (7) for the specific total receptor concentration, [T^*^], for which 50% of the receptors are dimeric and 50% are monomeric. For this particular concentration, [T^*^] = 2[*m*^*^] and *f*_*dimer*_ = *f*_*m*_ = 0.5 (where *f*_*m*_ is the fraction of monomers). At this particular concentration:

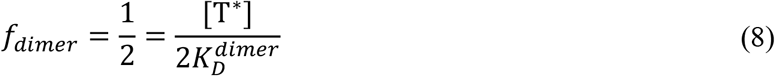

Therefore,

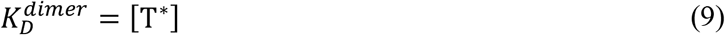

Eq. (9) is the condition under which Eq. (8) is satisfied. We thus see that when half of the receptors are in the dimeric state, the total receptor concentration, [T^*^], is equal to the dissociation constant, 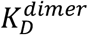. Unlike in the case of ligand binding to a protein discussed above, there are no assumptions, and this relation is always exact.

The meaning of the dissociation constant becomes more nebulous if the receptors associate into higher order oligomers. A monomer-oligomer model, given by the following reaction scheme, can be used to demonstrate this.

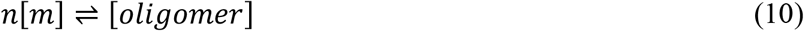

Where *n* is the order of oligomer and is greater than 2. The dissociation constant for this reaction can be defined as:

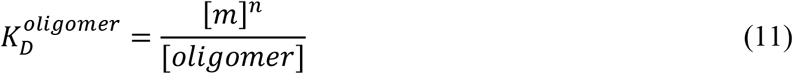

The units of *K*_*D*_ in Eq. (11) are not receptors per unit area but instead (receptors/μm^2^)^*n*^. Thus, the physical-chemical meaning of 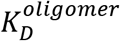 is not immediately obvious. In an analogy to the dimer case, we can write the oligomeric fraction as

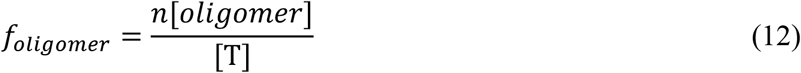

We solve equation (11) for the oligomer concentration and we substitute it into equation (12) to obtain.

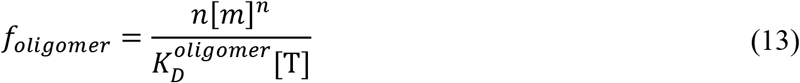

Now we write Eq. (13) for the specific receptor concentration, [T^*^] when [T^*^] = 2[*m*^*^] and *f*_*oligomer*_ = *f*_*m*_ = 0.5. This is the concentration at which half of the receptors are associated into oligomers and half are monomeric. For that particular concentration, the following equation holds.

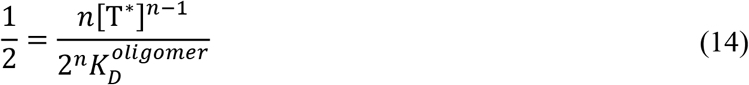

Now we solve Eq. (14) for [T^*^] and we define an effective dissociation constant as 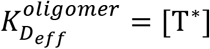.

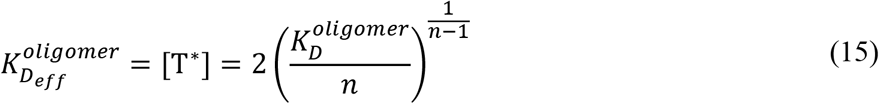

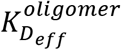 is thus the concentration for which 50% of the receptor is associated into oligomers and 50% is monomeric and has units of receptor concentration. Note that if we solve Eq. (15) for a dimer (*n* = 2), then we recover:

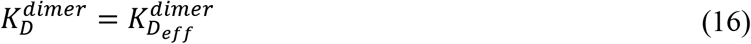

The effective dissociation constant, 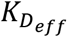, in Eq. (15) shares the same units of concentration and has a well defined physical meaning: They are given by the concentration for which 50% of the receptors are associated and 50% are monomeric, for each of the association models and for any *n*.

### FRET studies of EphA2 self-association in the plasma membrane

We used a quantitative FRET technique termed Fully Quantified Spectral Imaging FRET (FSI-FRET) (37), to study the self-association of full length EphA2 in the presence of the ligand ephrinA1-Fc. In these experiments the fluorescent proteins, eYFP or mTurquoise (a FRET pair), were attached to the C-terminus of the full length EphA2 with a flexible (GGS)_5_ linker. The images, acquired with a spectrally resolved 2-photon microscope, were analyzed to calculate (i) the apparent FRET efficiencies *E*_*app*_, (ii) the donor concentration, EphA2-mTurq, and (iii) the acceptor concentration EphA2-YFP in small areas of the plasma membrane of each cell(28). The data from 197 individual cells are shown in Figure 1. In Figure 1A we show the measured FRET efficiency, *E*_*app*_, as a function of EphA2-YFP concentration. We see that FRET increases as a function of concentration, in accordance with the law of mass action. Figure 1B shows the donor, EphA2-mTurq, concentration versus the acceptor, EphA2-YFP, concentration. Expression of the receptors varies over a significant concentration range since the receptors are introduced via transient transfection, yielding a binding curve.

**Figure 1.**
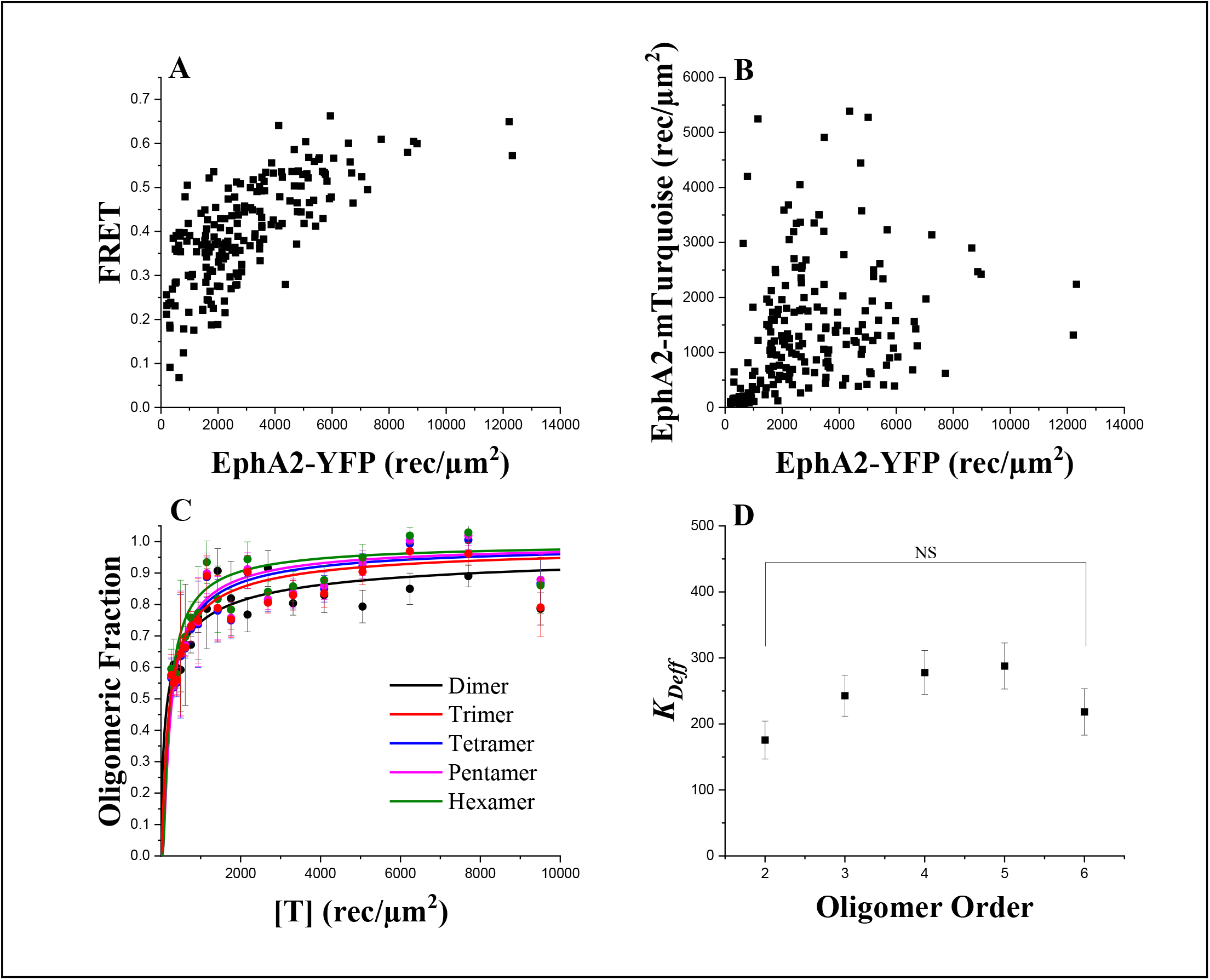
Determination of an effective dissociation constant from FRET data (A) Single cell FRET data for EphA2-mTuquoise (donor) and EphA2-YFP (acceptor), (B) EphA2-mTuquoise concentrations versus EphA2-YFP concentrations (C) Fits to oligomer models. (D) Comparison of EphA2 effective dissociation constants calculated for different oligomerization models. By ANOVA, there is no statistical significance between the values.

To analyze FRET data and to determine dissociation constants we follow the established FSI-FRET protocol (28). Specifically, we fit predictions for all models of association, from a dimer to a hexamer, given by Eq. (10) with *n* = 2 – 6, to the experimental data. The fitting procedure is described in detail elsewhere(28, 33). There are two unknown parameters in each fit (for an oligomer, or specific *n*): the dissociation constant 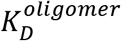 and 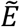, a structural parameter which depends on the relative distances and the dynamics of the fluorophores. For each association model (dimer through hexamer, *n* from 2 to 6) we write the oligomeric fraction as a function of 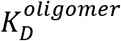 and the receptor concentration. We also calculate experimentally determined oligomeric fractions using equation (19) in Materials and Methods, after a correction for the so-called proximity FRET (33), as a function of 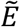. We vary the two unknown parameters, 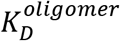 and 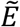, so the theoretical oligomerization curves provide the best fit to the experimentally measured values. In Figure 1 we show the theoretical oligomerization curves constructed for the best-fit 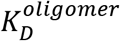 values. We also show the binned experimental oligomeric fractions, calculated for the best-fit 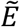 values.

In Table 1, we present the best-fit 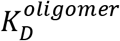 values. These values are very different and have different units. Furthermore, from N&B and FIF data we know that the distribution of EphA2 oligomer sizes is heterogeneous (29) with an average ∼4. None of the simple models of association given by equation (10) account for this heterogeneity. Thus, one might question the utility of the FRET method for studying EphA2 self-association.

**Table 1.** Dissociation constants calculated for different oligomerization models.

| Oligomer Model | $K_D^{oligomer}$ | $K_{D_{eff}}$ (rec/$\mu\text{m}^2$) |
| --- | --- | --- |
| Dimer | $180 \pm 30$ (rec/$\mu\text{m}^2$) | $180 \pm 30$ |
| Trimer | $40,000 \pm 10,000$ (rec<sup>2</sup>/$\mu\text{m}^4$) | $240 \pm 30$ |
| Tetramer | $1.1 \pm 0.4 \times 10^7$ (rec<sup>3</sup>/$\mu\text{m}^6$) | $280 \pm 30$ |
| Pentamer | $2 \pm 1 \times 10^9$ (rec<sup>4</sup>/$\mu\text{m}^8$) | $290 \pm 30$ |
| Hexamer | $2 \pm 2 \times 10^{11}$ (rec<sup>5</sup>/$\mu\text{m}^{10}$) | $220 \pm 40$ |

Using the best-fit 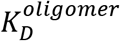 values in Table 1, we calculate effective dissociation constants using equation (10). They are also shown in Table 1 and they all have the same units, EphA2 concentration in the membrane. By ANOVA they are indistinguishable. Thus, all models used to fit the FRET data yield the same effective dissociation constant.

## DISCUSSION

It is known that EphA2, in response to its ligands, forms large oligomers in the plasma membrane that are heterogeneous in size(29, 38, 39). While EphA2 monomers are inactive, the EphA2 molecules in the oligomers activate each other by acting as both enzymes and substrates, attaching multiple phosphate groups to the intracellular tyrosines. Other RTKs, as well as other receptors, are also known to oligomerize, and the oligomer size often controls their function(31, 32). Previous work has shown that FRET experiments cannot discern the exact oligomer size(33). At best, they can differentiate between a dimer and a higher order oligomer, but a trimer cannot be distinguished from a tetramer or a higher order oligomer(33). Furthermore, the measured FRET for a heterogeneous population of oligomers will be the average of the FRET efficiencies for the different types of oligomers. Thus, it may appear to some that FRET measurements have significant limitations in characterizing the association state of membrane proteins.

Here we show that we can extract an effective equilibrium constant from an EphA2 FRET data set, no matter what model of association is assumed. Thus, the effective equilibrium constant can be determined even if the oligomer size is unknown. This equilibrium constant has a very well-defined physical meaning and biological significance. The effective dissociation constant is the EphA2 concentration for which half of the EphA2 molecules are monomeric and thus inactive, and the other half are associated into oligomers and are therefore active. We do not have to know the exact preferred oligomer size to draw this conclusion. Furthermore, we can say that for EphA2 concentrations below the effective dissociation constants, EphA2 will be predominantly monomeric and inactive. On the other hand, EphA2 will be predominantly associated into active dimers or oligomers once its concentration exceeds the effective dissociation constant.

This work therefore demonstrates the immense utility of FRET experiments in studies of receptor interactions even when the receptors form oligomers, not just dimers. It is highly recommended that FRET is used in conjunction with fluorescence fluctuation techniques, such as N&B(34), FIF(10, 29, 40), and PIE-FCCS(41-43), which directly report on the oligomer size but can present challenges in the calculations of dissociation constants.

## MATERIALS & METHODS

### Sample preparation

The EphA2 plasmid in the pcDNA3.1(+) vector was cloned in prior work (44). The plasmid encodes for human EphA2 tagged at the C-terminus with a fluorescent protein (either eYFP or mTurquoise) via a 15 amino acid GGS_5_ linker.(44)

HEK293T cells were purchased from American Type Culture Collection (Manassas, VA, USA). The cells were cultured in Dulbecco’s modified eagle medium (Gibco, #31600034) supplemented with 10% fetal bovine serum (HyClone, #SH30070.03), 20 mM D-Glucose and 18 mM sodium bicarbonate at 37 °C in a 5% CO_2_ environment. 24 hours prior to transfection, cells were seeded in 35 mm glass coverslip, collagen coated Petri dishes (MatTek, P35GCOL-1.5-14-C) at a density of 2.5*10^5^ cells per dish to reach ∼70% confluency at the day of the experiment. For transfection, Lipofectamine 3000 (Invitrogen, #L3000008) was used according to manufacturer’s protocol. Single transfections were performed using 1 – 3 ug plasmid DNA. Co-transfections were performed with 1 – 4 ug total plasmid DNA in a 1:3 donor:acceptor ratio. 12 hours after transfection the cells were rinsed twice with phenol-red free, serum free starvation media and then serum starved for at least 12 hours. The starvation media was supplemented with 0.1% BSA to coat the wall of the dishes.

### FRET Imaging

Before imaging, HEK 293T cells were subjected to reversible osmotic stress by replacing the serum-free media with a 37°C, 1:9 serum-free media:diH_2_O, 25 mM HEPES solution. In cells the plasma membrane is normally highly ruffled and its topology in microscope images is virtually unknown.(37) The reversible osmotic stress eliminates these wrinkles and allows the conversion of effective 3D protein concentrations into 2D receptor concentrations.(37) The swelling media was supplemented with 50 nM dimeric ephrinA1-Fc (R&D Systems, #602-A1-200). The cells were allowed to equilibrate for 10 minutes at room temperature. Images of cells were acquired using a two-photon microscope equipped with the OptiMiS spectral imaging system (Aurora Spectral Technologies). Two scans were performed for every cell– a FRET scan (λ_1_=840 nm) in which the donor (mTurquoise) is primarily excited and an acceptor scan (λ_2_=960 nm) in which the acceptor (YFP) is primarily excited. The output of each scan is composed of 300x440 pixels, where every pixel contains a full fluorescence spectrum in the range of 420 – 620 nm. Images of cells were acquired for up to 2 hours.

### FRET Data Analysis

The dependence of the FRET efficiencies on the fraction of receptors in the oligomeric state can be described using the “kinetic theory of FRET”(45):

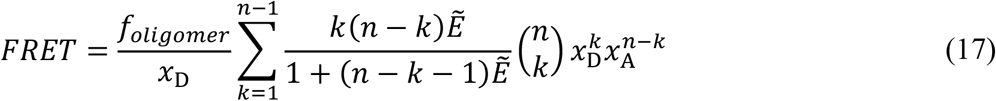

In Eq. (17), FRET is the FRET efficiency due to receptor oligomerization, which is calculated from the measured FRET after correction for the so-called “proximity FRET (see details for the correction in(33)). *f*_*oligomer*_ denotes the oligomeric fraction, *n* is the oligomer order, *x*_D_ & *x*_A_ are the fraction of donors and acceptors. The intrinsic FRET, 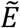, depends on the distance between the fluorophores, *r*, according to

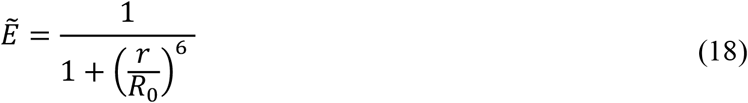

where Ro is the Forster radius. From (17), the oligomer fraction, *f*_*oligomer*_ is calculated as:

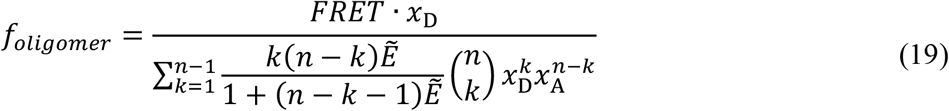

using the best-fit 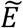. These are binned and shown in Figure 1 as symbols.

We also calculate the theoretical oligomeric fractions as a function of the dissociation constant, 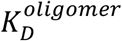, and the total concentration, [T] (solid lines in Figure 1). The derivation utilizes mass balance equations to re-write equation (10), and it published (28). Briefly, the mass balance equation for a dimer is:

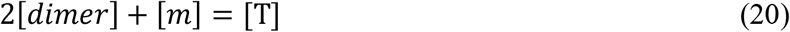

Then, Eq. (6) is solved for [*dimer*] and substituted into Eq. (20).

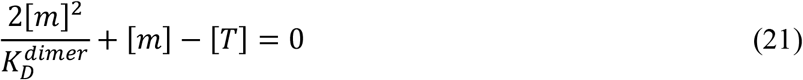

Eq. (21) is quadratic, and the solution is:

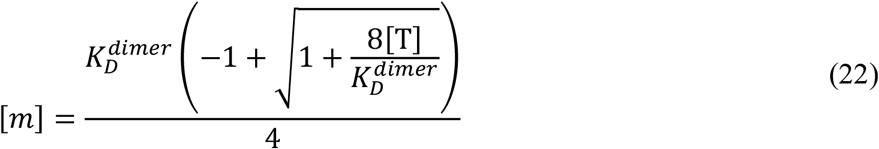

We substitute the values of [*m*] from (22) into Eq. (7) to calculate the dimeric fraction as a function of [T] and 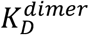, and we plot it as a solid line in Figure 1.

Similarly, for higher order oligomers the mass balance equation is given by:

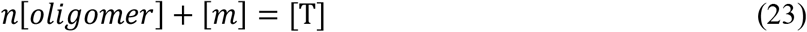

where *n* is the order of oligomer. This equation can be re-written as.

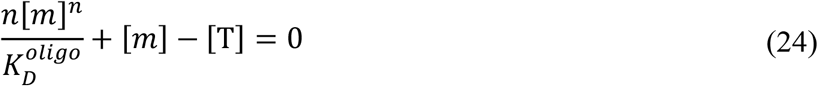

Just as in the dimer case, equation (24) is solved for [m] as a function of 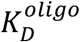 and [T], but now there is no analytical solution. Instead, the equation is solved numerically using MATLAB, and the oligomeric fraction are calculated by substituting the calculated [*m*] into eq. (13). They are plotted with solid lines in Figure 1.

